# Optimized cultivar deployment improves the efficiency and stability of sunflower crop production at national scale

**DOI:** 10.1101/2020.09.21.306076

**Authors:** Pierre Casadebaig, Arnaud Gauffreteau, Amélia Landré, Nicolas B. Langlade, Emmanuelle Mestries, Julien Sarron, Ronan Trépos, Patrick Vincourt, Philippe Debaeke

## Abstract

Plant breeding programs design new crop cultivars which, while developed for distinct populations of environments, are nevertheless grown over large areas during their time in the market. Over its cultivation area, the crop is exposed to highly diverse stress patterns caused by climatic uncertainty and multiple management options, which often leads to decreased expected crop performance.

In this study, we aim is to assess how finer spatial management of genetic resources could reduce the yield variance explained by genotype × environment interactions in a set of cropping environments and ultimately improve the efficiency and stability of crop production. We used modeling and simulation to predict the crop performance resulting from the interaction between cultivar growth and development, climate and soil conditions, and management practices. We designed a computational experiment that evaluated the performance of a collection of commercial sunflower cultivars in a realistic population of cropping conditions in France, built from extensive agricultural surveys. Distinct farming locations sharing similar simulated abiotic stress patterns were clustered together to specify environment types. We then used optimization methods to search for cultivars × environments combinations leading to increased yield expectations.

Results showed that a single cultivar choice adapted to the most frequent environment-type in the population is a robust strategy. However, the relevance of cultivar recommendations to specific locations was gradually increasing with the knowledge of pedo-climatic conditions. We argue that this approach while being operational on current genetic material could act synergistically with plant breeding as more diverse material could enable access to cultivars with distinctive traits, more adapted to specific conditions.

**Key message:** Crop simulation helps to analyze environmental impacts on crops and provides year-independent context information. This information is of major importance when deciding which cultivar to choose at sowing time.

## Introduction

### Context

The use of fertilizers, irrigation, and pesticides mitigated the effects of climatic hazards and had a large and positive impact on crop yield gains between 1960 and 2000 (Foley et al., 2005 ; Tilman et al., 2002). Currently, because of the simultaneous need to reduce inputs in agricultural systems, and the climatic uncertainty caused by climate change, the variability of cropping conditions increased as compared to late century conditions. Over its global cultivation area, a crop is thus exposed to highly diverse biotic and abiotic stress patterns, which can often lead to decreased crop performance regarding the expected level and cause yield gaps. Reducing these yield gaps could be achieved by two conflicting strategies: increasing inputs or increasing the crop resource-use efficiency (Sadras and Denison, 2016).

Plant breeding is a key asset to increase crop resource use efficiency by designing new cultivars adapted to distinct populations of environments (e.g. Voss-Fels et al., 2019 for wheat; Vear et al., 2003 for sunflower). Cultivars are nevertheless grown over large areas during their time in the market, which can encompass locations and management practices for which the cultivars were not designed for. In continental Europe, sunflower is mostly cultivated in eastern and southern regions (18.7 M hectares in 2018). In 2018, Russian Federation, Ukraine (together 49%, 26.9 Mt), and UE-28 (18%, 9.9 Mt) were the largest sunflower grain producers in the world accounting for 68% of global volume (FAO, 2020). In the last 30 years, the production area for the sunflower crop surged in Central and Eastern Europe (Ukraine, Russian Federation, + 9.6 M hectares together) while it decreased in Western Europe (France, Spain, - 1.1 M hectares together) (FAO, 2020). These changes in global acreage, while accounted for in plant breeding programs can lead to sub-optimal use of the developed genetic resources.

*Phenotypic plasticity* is defined as the range of phenotypes a single genotype can express as a function of its environment (Nicotra et al., 2010). At the population level, this process might result in the relative variation in the performance of cultivars across environments. In this case, it is referred to as genotype × environment interactions in plant breeding and agricultural extension (Van Eeuwijk et al., 2016), which can outweigh the main genotype effect and can explain up to 20% of total yield variance observed in multi-environment trials (e.g. Foucteau et al., 2001, for sunflower). Consequently, yield gap reduction is limited by our ability to identify favourable combinations of cultivars and cropping conditions, given the resources available to experiment among possible combinations in the target population of environments (Comstock, 1976; Hammer and Jordan, 2007). Crop improvement can be viewed as a search strategy on a genotype-environment space (Hammer et al., 2006) to manage the genetic and environmental resources more efficiently by taking advantage of phenotypic plasticity.

Plant modeling approaches have emerged as a method to complement and improve the resource-limited experimental exploration of the adaptation landscape (Chapman et al., 2003; Messina et al., 2006; Hammer et al., 2006; Messina et al., 2011). Such models represent biological processes linked to plant growth and development as a function of time, environment (climate, soil, and management), and genetic diversity. Crop simulation models have the capacity to explore consequences of potential agronomic and breeding interventions in the design of crops for production systems (Sinclair et al., 2019; Hammer et al., 2019) and have been successfully used in many cases (see Chenu et al., 2017 for a review in wheat).

Two cases are particularly relevant to leverage phenotypic plasticity and support cultivar deployment strategies.

The first one, referred to as *environmental characterization* or *envirotyping* (see Xu, 2016 for a review), aims to analyze environmental impacts on crops and produce a set of categories (environment-types) grouping multiple cropping conditions (e.g. distinct locations and years) into comparable stress scenarios. While climate, soil, and management data could directly be used as classifiers, simulation was successfully used to generate new variables representing abiotic stress *per se* by accounting for the interactions between the plants and their environments (Chenu et al., 2013), potentially more informative than raw environmental data (Mangin et al., 2017). These variables, such as time series of water supply:demand ratio, can then be clustered into distinct stress scenarios over a large spatial and temporal scale (Chenu et al., 2013; Gosseau et al., 2019).

In the second case, crop models are at the core of numerical experiments, aiming to complement and extend field experiments such as multi-environment trials (MET). The capacity to sample the target population of environments for a set of cultivars along with the possibility to formalize optimization problems enabled applications in cultivar selection (e.g. Li et al., 2013), trait evaluation in multiple environment types (Chapman et al., 2002), or trait optimization for performance or resistance criteria (Semenov and Stratonovitch, 2013; Paleari et al., 2015; Picheny et al., 2017a).

### Problem and aim

In France, the commercial sunflower cultivars are a subset of those developed for the European market, while the French cropping conditions do not necessarily reflect those for which European cultivars were developed (Debaeke et al., 2017; Gosseau et al., 2019). In this study, we aim to assess how finer spatial management of available genetic resources could reduce the yield variance explained by genotype × environment interactions in actual cropping conditions and ultimately improve the efficiency and stability of crop production.

For that, we developed an approach (outlined in Figure 1) where we used postal surveys to describe the cultivation area and crop management practices at the farm scale, over the national acreage. We then designed a realistic numerical experiment based on this information, as a factorial combination of distinct farm locations (including crop management) and a collection of commercial cultivars. High-resolution climate and soil gridded datasets were used to complete survey data and enhance climatic coverage. Simulation allowed to predict the crop performance resulting from the interaction between cultivar growth and development, climate and soil conditions, and management practices. Finally, we defined and solved optimization problems to assess different cultivar deployment strategies leading to increased yield expectations.

**Figure 1.**
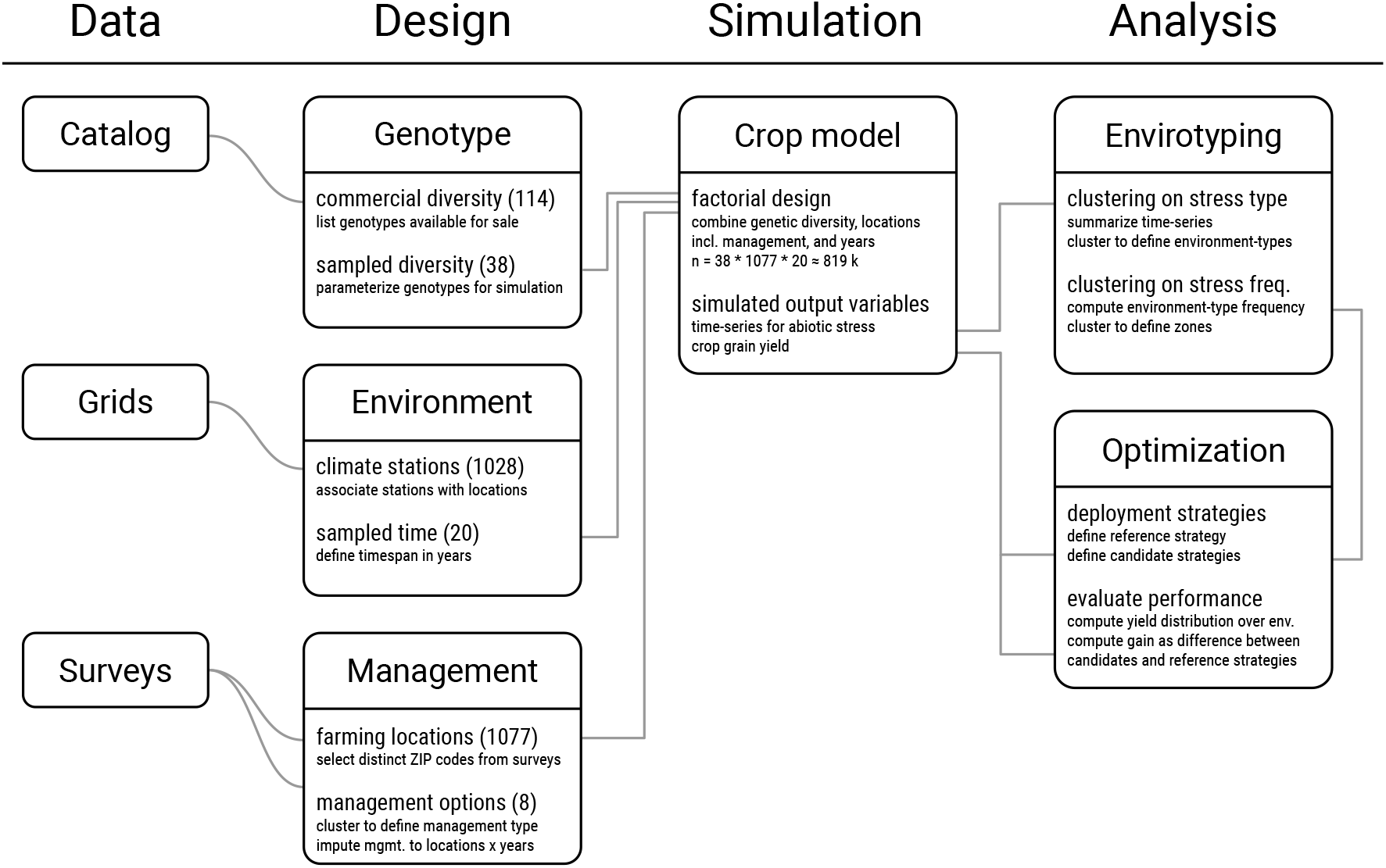
Outline of the approach used to compare cultivar deployment strategies. A synthetic view of the design, simulation, and analysis steps (columns). In the *design* step, each box presents the number of modalities used for genotype, locations, climate, and management factors along with a summary of the method used. In the *simulation* step, the box summarizes the design of the numerical experiment along with simulated variables. The *analysis* step illustrates how the envirotyping and optimization methods are linked together and how yield gains are computed.

## Material and methods

### Farm data

Farming data concerning the sunflower crop in France were collected from extensive postal surveys conducted by an agricultural extension institute (*Terres Inovia*). Between 1997 and 2013, 11 years of data were gathered, representing 11969 farming situations in total (from 1077 distinct locations). In total, 14 crop management variables related to the sunflower crop were collected, including qualitative (e.g. cultivar, fungicide application, soil depth class) and quantitative variables (e.g. sowing date and density, nitrogen fertilization amount). We focused the study on variables linked to abiotic stress (water, radiation, temperature, nitrogen).

### Description of farming locations

The farm data featured the ZIP code, which was used as an approximation of the actual farm geographical location (not collected).

We used the European Soil Database derived data (Hiederer, 2013, http://esdac.jrc.ec.europa.eu/content/esdb-derived-data, 1×1 km grid cells) to obtain the quantitative soil properties corresponding to each location, i.e. total available water content, soil depth available to roots, bulk density, coarse fragments for two soil layers ([0, 30 cm],] 30 cm, rooting depth]).

20 years (1994-2013) of climatic data were obtained from the French national center for meteorological research (CNRM), using an 8×8 km grid dataset based on the interpolation of observed climatic data (Quintana-Segui et al., 2008, Meteo-France SAFRAN). Farm locations were associated with the closest climate grid point. Five climatic variables were used: minimal and maximal temperature (°C), evapotranspiration (mm), global radiation (MJ m^-2^), and precipitation (mm).

### Cluster analysis of management practices

To reduce the number of modalities in surveyed management practices, we used a classification analysis to cluster farming situations that shared similar management practices. First, factor analysis for mixed data (FAMD, Pagès, 2004) was performed on the data to simplify variables that are correlated. We then performed a hierarchical cluster analysis (HCA, Kaufman and Rousseeuw, 2009) on the variables resulting from FAMD to cluster management practices.

### Crop modeling and simulation

#### Model description

We used crop modeling and simulation to predict the grain yield in non-observed field environments. SUNFLO is a process-based simulation model for sunflower that was developed to simulate grain yield and oil concentration as a function of time, environment (soil and climate), management practices, and genetic diversity (Casadebaig et al., 2011; Lecoeur et al., 2011). Predictions with the model are restricted to potential or attainable yield (Van Ittersum and Rabbinge, 1997): only the main limiting abiotic factors (temperature, radiation, water, and nitrogen) are included in the algorithm. Weeds, pests, and diseases are therefore not accounted for in this study. The model simulates the main soil and plant functions: root growth, soil water and nitrogen dynamics, plant transpiration and nitrogen uptake, leaf expansion and senescence, and biomass accumulation. Four climatic variables are used as daily inputs for simulation: mean air temperature (°C, 2m height), global incident radiation (MJ.m^-2^), potential evapotranspiration (mm, Penman–Monteith) and precipitation (mm). Soil properties are defined by its texture, maximum depth and daily nitrogen mineralization potential, while initial soil conditions are defined by residual mineral nitrogen and initial water content. The needed management practices include sowing date and density, irrigation and nitrogen fertilization (timing and amount). Globally, the SUNFLO crop model has about 50 equations and 64 parameters split into 33 species-dependent, 10 genotype-dependent, and 21 environment-related parameters. A report that summarizes the equations and parameters used in the model is available as supplementary information in Picheny et al. (2017a).

SUNFLO was evaluated both on specific research trials (40 trials, 110 cultivar × environment combinations) and on agricultural extension trials that were representative of its targeted use (96 trials, 888 cultivar × environment combinations). Over these two datasets, the model was able to simulate significant G × E interactions and rank genotypes (Casadebaig et al., 2011, 2016). From these evaluations, we considered that SUNFLO was accurate enough to support optimization methods, i.e. allows discrimination between two given cultivars.

#### Experimental design and numerical experimentation

To describe a realistic target population of environments for sunflower cropping in France, we used a factorial design crossing a cultivar list of commercial sunflower hybrids (*g* = 38), the surveyed farming locations (soil and crop management, *e* = 1077) and 20 years of historical climate data (1994-2013, *t* = 20). The resulting design represented 818520 virtual fields (*n* = *g* × *e* × *t*).

Genetic diversity was represented with *g* = 38 cultivars resulting from the intersection of the list of cultivars actually available for sale in 2017 (n=125, Terres Inovia, 2017) and those whose morphological and physiological characteristics were assessed for simulation (n=69). Such characteristics were necessary to simulate genotype × environment interaction with the SUNFLO crop model, which uses measured phenotypic traits as genotype-dependent parameters. The estimation of these parameters is based on plant phenotyping, with methodology and protocols fully described for field (Casadebaig et al., 2011, 2016) and controlled conditions (Casadebaig et al., 2008; Gosseau et al., 2019).

Briefly, each cultivar is currently described by 10 parameters, which can be organized in four groups: phenology (2 parameters), leaf architecture (4), response to water constraint (2), and biomass allocation to the grains (2) (Table 1). Furthermore, we assumed that this set of 10 traits were sufficient to describe the adaptation of a variety to a location. Other traits, not accounted for in the crop model can also drive plant response to environment (such limitations are further discussed in Casadebaig et al., 2011).

**Table 1:** Description and variation range for genotype-dependent parameters of the SUNFLO model. The maximum and minimum values reported represent the genotypic variability observed among phenotyped sunflower cultivars. Phenological traits were measured on the whole microplot (50% of the population required to reach the stage). Architectural traits were measured of 5 plants per microplot. Potential harvest index was estimated on 10 plants per plot, as the ratio of seed to shoot biomass, including senescent organs. Potential seed oil content was determined as the 9th decile of the distribution of oil concentration values measured in national networks for cultivar evaluation. Protocols for trait measurement are detailed in Casadebaig et al. (2016).

| Function | Description | Unit | Min | Max |
| --- | --- | --- | --- | --- |
| Phenology | Temperature sum from emergence to the beginning of flowering | *Cd* | 744.25 | 906.60 |
| Phenology | Temperature sum from emergence to seed physiological maturity | *Cd* | 1460.80 | 2054.90 |
| Architecture | Potential number of leaves at flowering | *leaf* | 22.20 | 36.67 |
| Architecture | Potential rank of the plant largest leaf at flowering | *leaf* | 12.30 | 23.20 |
| Architecture | Potential area of the plant largest leaf at flowering | *cm<sup>-2</sup>* | 139.49 | 670.00 |
| Architecture | Light extinction coefficient during vegetative growth | - | 0.78 | 1.00 |
| Response | Threshold for leaf expansion response to water stress | - | -15.57 | -2.15 |
| Response | Threshold for stomatal conductance response to water stress | - | -14.21 | -5.51 |
| Allocation | Potential harvest index | - | 0.25 | 0.51 |
| Allocation | Potential seed oil content | % | 47.77 | 62.30 |

Surveyed management practices were depicted with eight management systems resulting from the cluster analysis. These management systems were used for the simulation analysis and qualified by the sowing date, plant density, and the amount of mineral nitrogen applied. (modalities indicated in Table 3). In sunflower, nitrogen fertilization is mainly applied at sowing or during early growth: one single application of 45 kg/ha of mineral nitrogen is the dominant practice (Enquêtes Pratiques Culturales, AGRESTE, 2007). Because the survey data did not contain the timing of nitrogen application, we used the sowing date as the application date in simulations. Less than 5% of the area cultivated with sunflower in France is currently irrigated (Enquêtes Pratiques Culturales, AGRESTE, 2007), we, therefore, did not consider irrigation as a management option to alleviate water deficit in simulations.

As some farms with different management systems shared the same postal code, the most frequent management system for each location-year combination was selected to assign a single management system to each location-year combination. Because the surveys were not conducted every year, the management systems missing from the factorial design were imputed assuming that management on the location did not change until the next surveyed date (last observation carried forward, LOCF).

For each genotype x location x year combination, five output variables were simulated by the SUNFLO crop model: seed yield at harvest, and four time-series corresponding to the impact of abiotic stressors on crop photosynthesis, i.e. heat stress as a function of mean air temperature, cold stress as a function of mean air temperature, water stress as a function of water deficit (simulated with the fraction of transpirable soil water, FTSW) and nitrogen stress as a function of nitrogen deficit (simulated by the nitrogen nutrition index, NNI, Debaeke et al., 2012).

Only in a few cases, the SUNFLO crop model failed to simulate seed production (∼ 4.5% of genotype x environment combinations) either because of excessive water deficit or the predicted maturity date was too late in the season for a realistic harvest date. One cultivar was discarded from the analysis because simulated yield values were outliers in all environments. The difference between modalities in studied factors (locations, genotypes) for each step of the study is reported in Table 2.

**Table 2.**
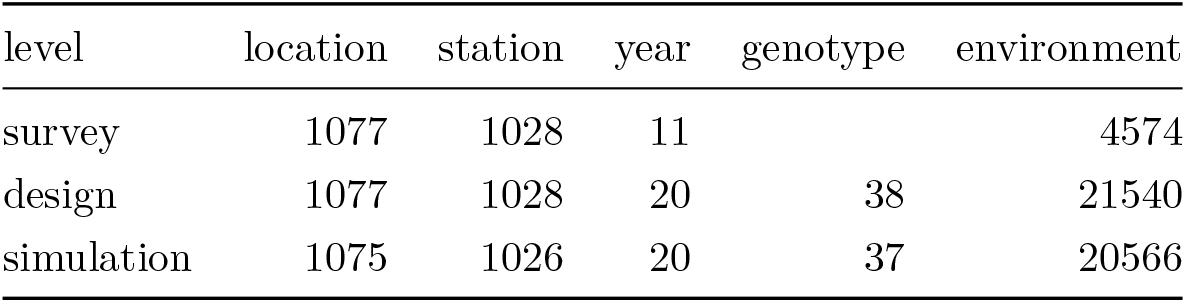
Sampled cropping conditions at the survey, numerical design, and simulation levels. The modalities of the sampled factors are reported at different steps of the study: postal survey, numerical design of experiments, and actual simulated results. Columns correspond to the number of distinct farm geographical locations (*location*), the distinct grid points used as climate data source (*station*), the distinct climatic years and genotypes (survey data did not include the genotype name, only the maturity group), and environments defined as the distinct combinations of locations and years.

| level | location | station | year | genotype | environment |
| --- | --- | --- | --- | --- | --- |
| survey | 1077 | 1028 | 11 |  | 4574 |
| design | 1077 | 1028 | 20 | 38 | 21540 |
| simulation | 1075 | 1026 | 20 | 37 | 20566 |

**Table 3.** Classification of management practices for sunflower in France. Eight management types (named M1-M8) resulted from the cluster analysis of agricultural survey results (14 management-related variables, 11k observations), with their proportion indicated in percent (*p*). Six variables were used to characterize the management type: sowing date (dd/mm), crop density (plants.m^-2^), mineral nitrogen fertilization (kg.ha^-1^), frequency of boron application (%), frequency of fungicide application (%) and frequency of early cultivar use (%). Label was assigned *a posteriori*, based on the interpretation of those six variables.

| group | label | p | sowing | density | nitrogen | boron | fungicide | precocity |
| --- | --- | --- | --- | --- | --- | --- | --- | --- |
| M1 | Low chemical inputs | 13 | 20/04 | 6 | 15 | 19 | 5 | 64 |
| M2 | Early cultivar use | 14 | 15/04 | 6 | 46 | 31 | 1 | 89 |
| M3 | Early sowings | 13 | 20/04 | 6 | 45 | 12 | 6 | 72 |
| M4 | Early sowings, boron | 8 | 05/04 | 6 | 50 | 85 | 36 | 80 |
| M5 | Very late sowings, very early cultivars | 12 | 10/05 | 6 | 38 | 13 | 5 | 66 |
| M6 | Late cultivars | 15 | 15/04 | 5 | 53 | 13 | 7 | 9 |
| M7 | Late sowings, early cultivars | 11 | 01/05 | 6 | 62 | 22 | 11 | 68 |
| M8 | High chemical inputs | 14 | 20/04 | 5 | 60 | 82 | 82 | 0 |

### Clustering of environments

Each cropping condition (location-year combination), referred as *environment*, was characterized by four indices that were computed by integrating the simulated time series of abiotic stressors (*X*_*te*_) over the crop cycle (*d* days between sowing and harvest) and averaging for genetic diversity (*n* genotypes) (equation 1).

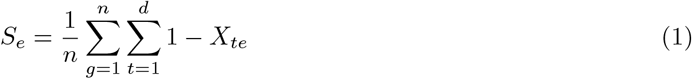

with *S*_*e*_, level of abiotic stressor *e*, i.e. for cold (LTRUE), heat (HTRUE), water (WRUE) or nitrogen (NRUE) stress. *X*_*te*_, value of daily simulated impact of the abiotic stressor *e* on crop photosynthesis.

We then proceeded with two successive hierarchical cluster analysis, based on Euclidian distances and Ward’s variance clustering method. The final cluster number was defined according to the between:total sum of square ratio and ecophysiological interpretability. The first step of clustering was applied on the matrix of scaled abiotic stress index, to identify groups of environments with similar abiotic stress patterns, referred to as environment-types (e.g. see Chenu et al., 2013 for water-deficit patterns). For the sunflower crop, this method was also used in previous studies about environmental characterization (Mangin et al., 2017; Gosseau et al., 2019). The second step of clustering was applied on the frequency of environment types per location, to identify groups of geographical locations with a similar frequency of environment types, independently of climatic variability, referred as agricultural zones (Beillouin et al., 2018).

### Baseline and optimization for cultivar deployment strategies

Here, our aim was to evaluate the impact of different cultivar deployment strategies on the distribution of grain yield across the population of cropping conditions. For that, we compared baseline with optimized deployment strategies (Table 4).

**Table 4.** Summary table of the comparison between reference and tested cultivar deployment strategies. The cultivar deployment strategies (column *strategy*) were characterized by the number of decisions taken to adapt cultivars to the cropping conditions (*decision*), the number of distinct cultivars that were actually chosen (*diversity*), the distance between the reference and each optimized strategy (*distance*, in q.ha^-1^), the median value of the yield gain at the population level (*gain_tpe*, in %), the expected shortfall gain at the population level (*es_tpe*, in %) and at the farm level (over the 20 sampled years, *es_farm*, in %). Expected shortfall is a risk measure focusing on the less profitable outcomes (here, the mean gain for the 10% worst cases).

| strategy | decision | diversity | distance | gain_tpe | es_tpe | es_farm |
| --- | --- | --- | --- | --- | --- | --- |
| random | 20566 | 37 | 3.4 | -5.7 | -5.8 | -10.2 |
| global | 1 | 1 | 2.2 | 7.0 | 4.0 | 6.1 |
| env_zone | 4 | 2 | 2.5 | 7.8 | 12.5 | 7.5 |
| farm | 1075 | 18 | 2.6 | 7.9 | 12.7 | 7.5 |
| farm_year | 20566 | 29 | 2.8 | 8.5 | 13.3 | 9.4 |

Two baseline deployment strategies were defined: *random*, a random choice of cultivar per cropping condition, and *reference*, based on surveyed cultivar acreage data. The *random* strategy was only used as a control and the more realistic *reference* strategy was the one used as a reference for the comparison with optimized strategies. We defined the *reference* strategy to represent the current cultivar recommendations and farmers’ choices. In this strategy, we assumed that the best cultivars were those that had a major commercial success. This list was obtained by selecting *s* distinct cultivars among the five most-grown cultivars (based on the actual sown area) and for each of the last five years of the study (2009-2013). The reference yield (*Y*_0_) for each cropping condition under this strategy was the average of this cultivar list, i.e. 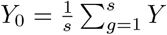, with *Y*, simulated grain yield, and *g*, genotypes from the cultivated list.

We defined four optimization strategies to recommended cultivars, ranging from a global adaptation strategy to a local one. The resolution of strategies differed by the number of decisions (cultivar choice) made according to the level of knowledge about the cropping conditions:

1. *global*, one choice for all the population of environments;
2. *zone*, one choice per agricultural zone (4 choices, see the previous section);
3. *farm*, one choice per farm location (1075 choices);
4. *farm_year*, one choice per farm:year combination (20566 choices).

Using the *zone* strategy as an illustration, the optimization algorithm first computes the mean values by cultivars for each agricultural zones (*Z1, Z2, Z3, Z4*), and the best cultivar is then selected for each zone, thus providing a list of optimized choices (4 cultivars, potentially distinct). This list is then used to obtain the optimized yield (*Y*_*m*_) distribution, by filtering all simulated cultivar × environments on optimal cultivar choices. The relative yield gain distribution was computed as 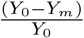. The performance of optimization strategies was estimated by computing the median value of the relative yield gain distribution. The distance between strategies was computed with the rooted-mean-squared differences between yield gain distributions. We evaluated yield stability in a conservative way, focusing on the less profitable outcomes by computing the mean of yield values below the 0.1 quantile value (expected shortfall, Acerbi and Tasche, 2002) of the yield gain distribution. The Expected Shortfall was computed for the yield distribution at the population of environment level and at the farm level, over the sampled years.

To avoid using all the data to select and evaluate the performance of strategies, we used a k-fold cross-validation resampling strategy (with folds corresponding to the year factor) to select optimal cultivars on a partition of the data (all years minus one) and estimate the performance of these choices on the other partition (one year).

### Software and data analysis

All data processing, statistical analysis and visualization were performed with the R software version 4.0.2 (R Core Team, 2018) with additional R packages *dplyr* (data processing, Wickham et al., 2018), *ggplot2* (visualization, Wickham, 2016), rsample (resampling, Kuhn et al., 2019), and *knitr* (reporting, Xie, 2015). Both Factor Analysis For Mixed Data (FAMD) and Hierarchical Cluster Analysis (HCA) were performed with the *FactoMineR* R package (Lê et al., 2008). The source code for the SUNFLO simulation model is available on INRA software repository [https://forgemia.inra.fr/record/sunflo.git]. The INRA VLE-RECORD software environment (Quesnel et al., 2009; Bergez et al., 2013) was used as simulation platform.

## Results

### Cropping conditions for the sunflower crop in France

The surveyed farms were globally matching the main regions of sunflower production (Figure 2), with sampled regions (i.e. containing sampling points) representing over 95% of the total cumulated acreage (14.9 Mha over 1994-2013).

**Figure 2.**
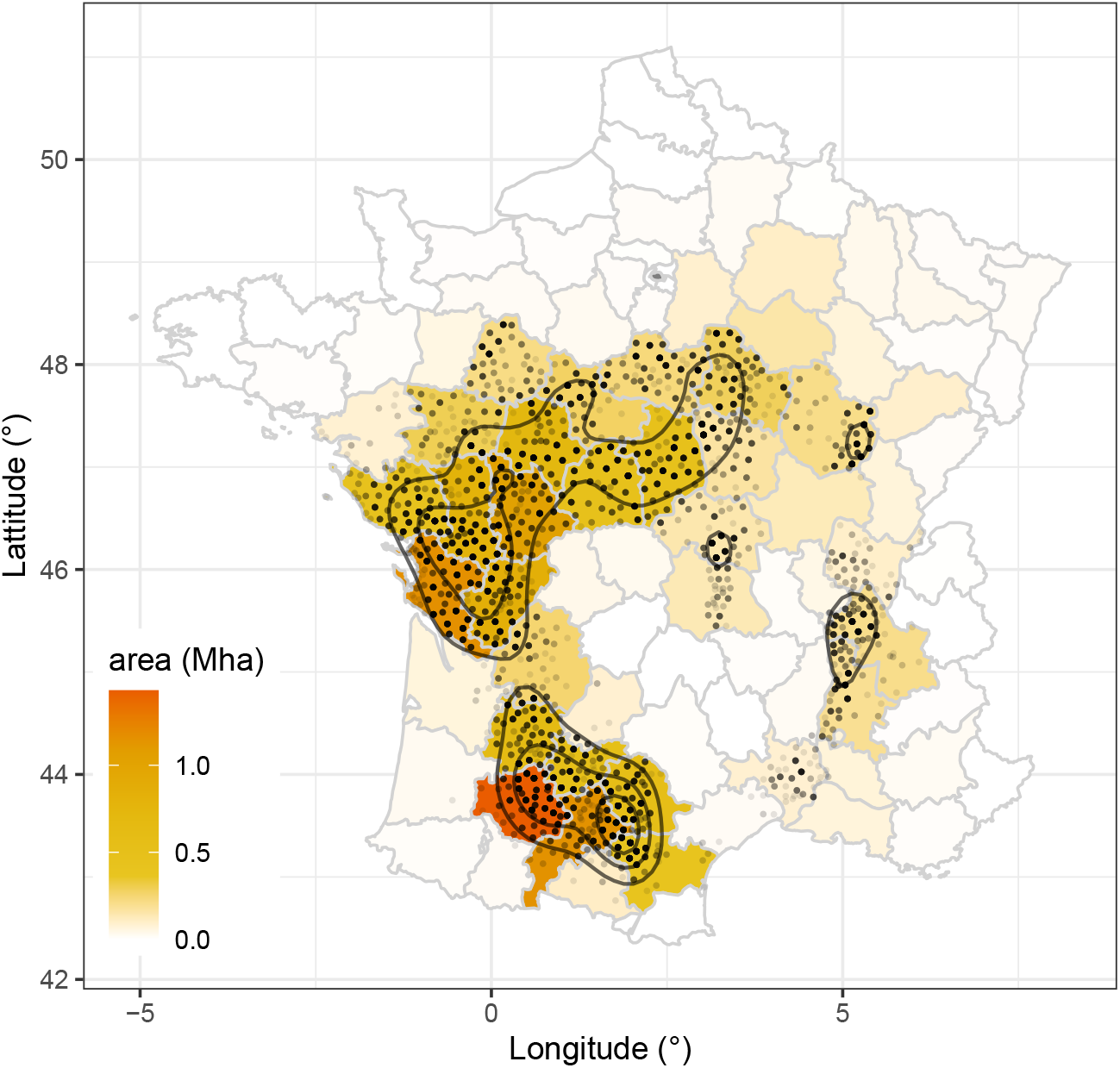
Map of sampled locations in the sunflower farm-level surveys. Points are the location of the 1077 unique farms sampled over 11 years (source: Terres Inovia surveys) with contour curves approximating the point density. Colors indicate the cumulated cropped area (Mha) on each French sub-regions (*départements*) for sampled years (1994-2013).

The classification analysis on management practices resulted in 8 clusters (labeled from M1 to M8 in Table 3) common to the 11 years and the five regions of survey. Clusters were roughly balanced with average proportions between 8 and 15% of farmers. Among the 14 variables of the survey, the amount and type (organic/mineral) of nitrogen fertilization, fungicide use, and the precocity of cultivars were the main discriminative variables acting on clusters.

M1 management type could be considered as low input (lowest nitrogen fertilization, low fungicide use). On the opposite, M8 type could be considered as high input management: farmers systematically use a fungicide with high nitrogen fertilization and frequent boron application. Other management modalities were less discriminated by input levels and were differing mostly by the crop sowing dates (early implantation for M4 or very late for M5), and cultivar precocity (e.g. early cultivar to compensate for late implantation in M5).

Regarding the cultivated genetic diversity, we observed a strong decrease in the cumulated proportion of area for the five most sown cultivars, going from 47% to 18% (rate was -2.5% per year, p=1.2e-7, R^2^=0.89).

### Environmental characterization

#### Climatic and soil variability

Globally, a large proportion of the sunflower cropping area is exposed to climatic water deficit, particularly in the South-West and West regions (Figures 2 and 3A).

**Figure 3.**
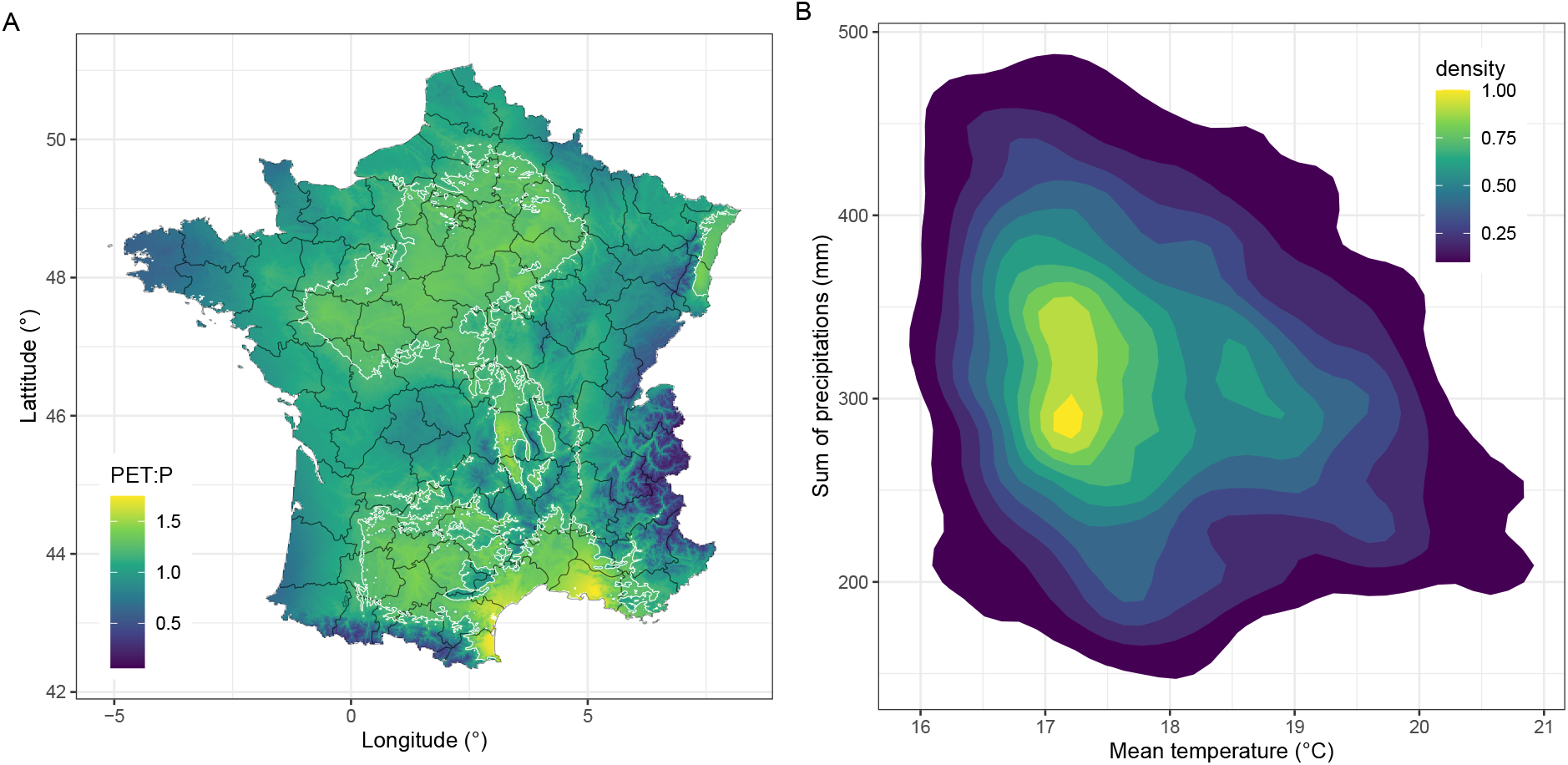
Climatic variability in the sampled cropping conditions. The left panel (A) displays a map of potential evapotranspiration to precipitation ratio (1 / aridity index, data from Zomer et al. (2008)). The white contour line corresponds to a PET:P ratio of 1.2, indicating geographical locations prone to water deficit. The right panel (B) shows climatic variability in the sampled gridded points (1028) over 20 years (n=20560). Color is the 2D density estimate of the sum of precipitations as a function of the mean temperature for the population of environments. Mean and sum were computed between average sowing and harvest dates (April 15th to September 15th).

For the considered climatic conditions (20560 stations x year population), weather records during the cropping season (Figure 3B) indicated that the mean sum of rainfall was 319 ± 94 mm, ranging from 80mm to 993mm. Mean temperature was 17.9 °C, ranging from 9.9 °C to 23.8 °C. In this population of environments, 1.6% corresponds to semi-arid conditions (aridity index ∈]0.2; 0.5]) and 8.6% to dry sub-humid (aridity index ∈] 0.5; 0.65]), according to Middleton et al. (1997) boundaries for dryland classification. Mean soil water capacity in the population of environments was 141 mm, ranging from 38 mm to 350 mm (*sd* of 51 mm)

#### Simulated abiotic variables allowed to define coherent environment types

Figure 4 displays the variability of the simulated abiotic stressors as a function of environment type resulting from the clustering of the population of environments. Cutting the dendrogram to four groups gave a ratio between:total sum of square of 63 % and allowed a sensible interpretation of stress patterns. Stress patterns were contrasted between groups, allowing to comment and name environment types (left to right in Figure 3) based on their distributions. The first group (*optimal*) was characterized by comparatively low mean stress levels corresponding to near-optimal cropping conditions. The second group (*cold*) displayed a high level of cold stress, while the other stresses were at a very low level. The third group (*heat and drought*), was the only group with an important level of heat stress which was associated to moderate level of water deficit. The fourth group (*drought and nitrogen*) was characterized by unfavorable cropping conditions, with the strongest level of water deficit associated with nitrogen deficit.

**Figure 4.**
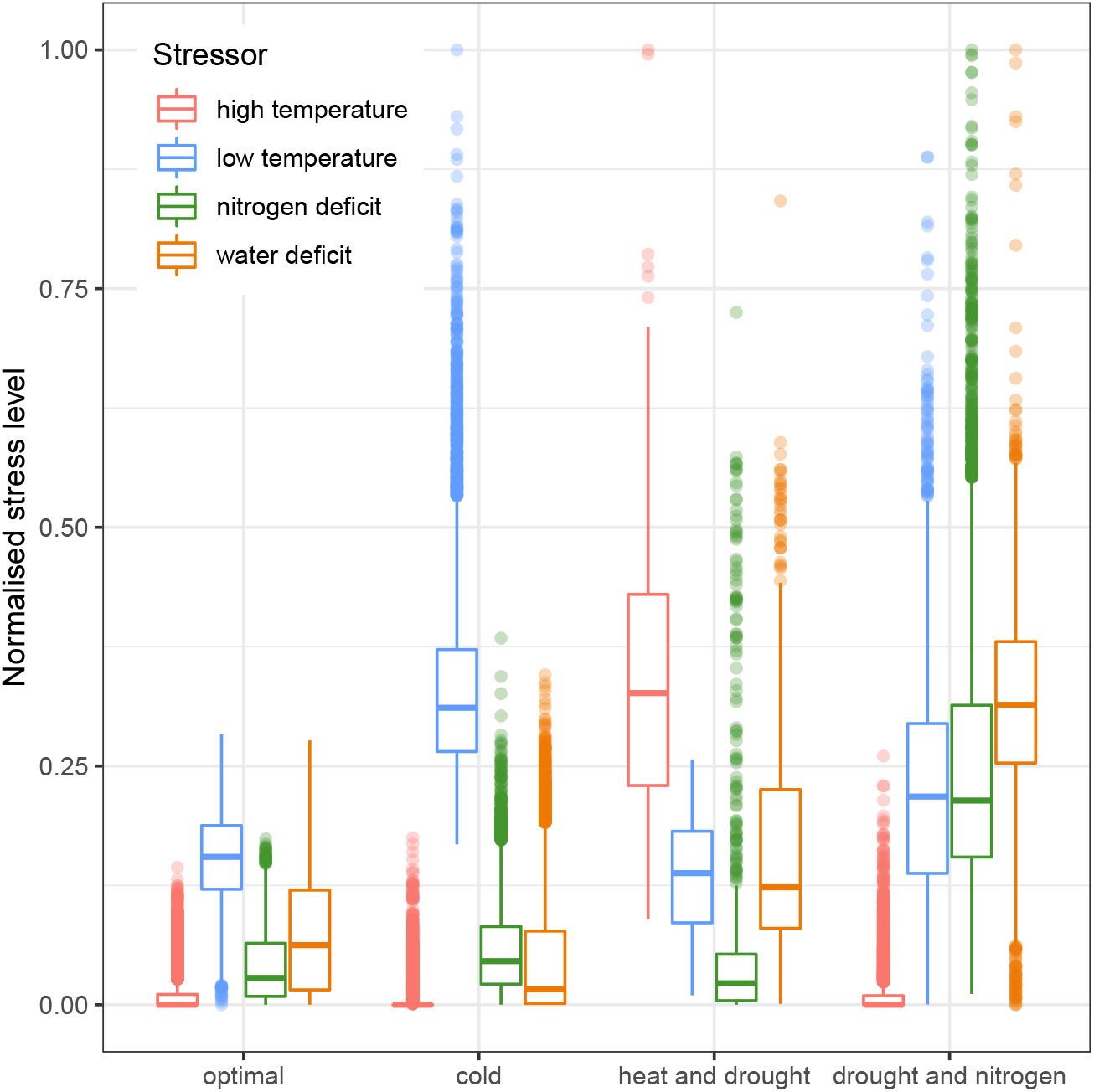
Distribution of abiotic stress levels as a function of defined environment types. Simulation was used to characterize the seasonal abiotic stress patterns at the crop level. Each environment was characterized by a vector of four abiotic indices, i.e. stressors integrated over the cropping season (high and low temperatures, nitrogen, and water deficit impact on crop photosynthesis). Hierarchical clustering was used to group abiotic patterns into four environment types, *a posteriori* named after the observed stress pattern. Values of abiotic stressors were rescaled (unity-based normalization) to allow comparisons between stressors.

#### Temporal and spatial distributions of environment types are highly variable

The first cluster analysis allowed to assign a distinct type of environment to each location for each year (1075 locations × 20 years, 20566 cropping environments). To better understand how these environment types were related to actual cropping conditions, we displayed the evolution of the proportion of each environment type as a function of time (Figure 5A) and the succession of environment types for each geographical location through years (Figure 5, panels C-F). At the national scale, *cold* and *optimal* environment types were the most frequent (average frequency of 51% and 28% respectively), followed by *drought and nitrogen* type (17%). The frequency of occurrence of environment types was varying strongly with time, with some peculiar years such as 2003 (very high proportion of *heat and drought* type), 2007 and 2011 (high cumulated proportion of environment types with low water deficit), and 2012 (the highest proportion of unfavorable *drought and nitrogen* environment type). From a geographical point of view (Figure 5, panels C-F), the location of *optimal* environment types matched the area with the most sunflower acreage (South-West France). In contrast, *drought and nitrogen* environments were not specifically bound to climate and could be located anywhere in the cropping area (e.g. panel F), with however some recurring locations (South-East). Even if the frequency of environment types could be related to national yield for specific years (low national yield in 2003 associated with high-temperature stress and record years in 2007 and 2011), we did not find a strong relationship between the frequency of environment types and the national yield (Figure 5B).

**Figure 5.**
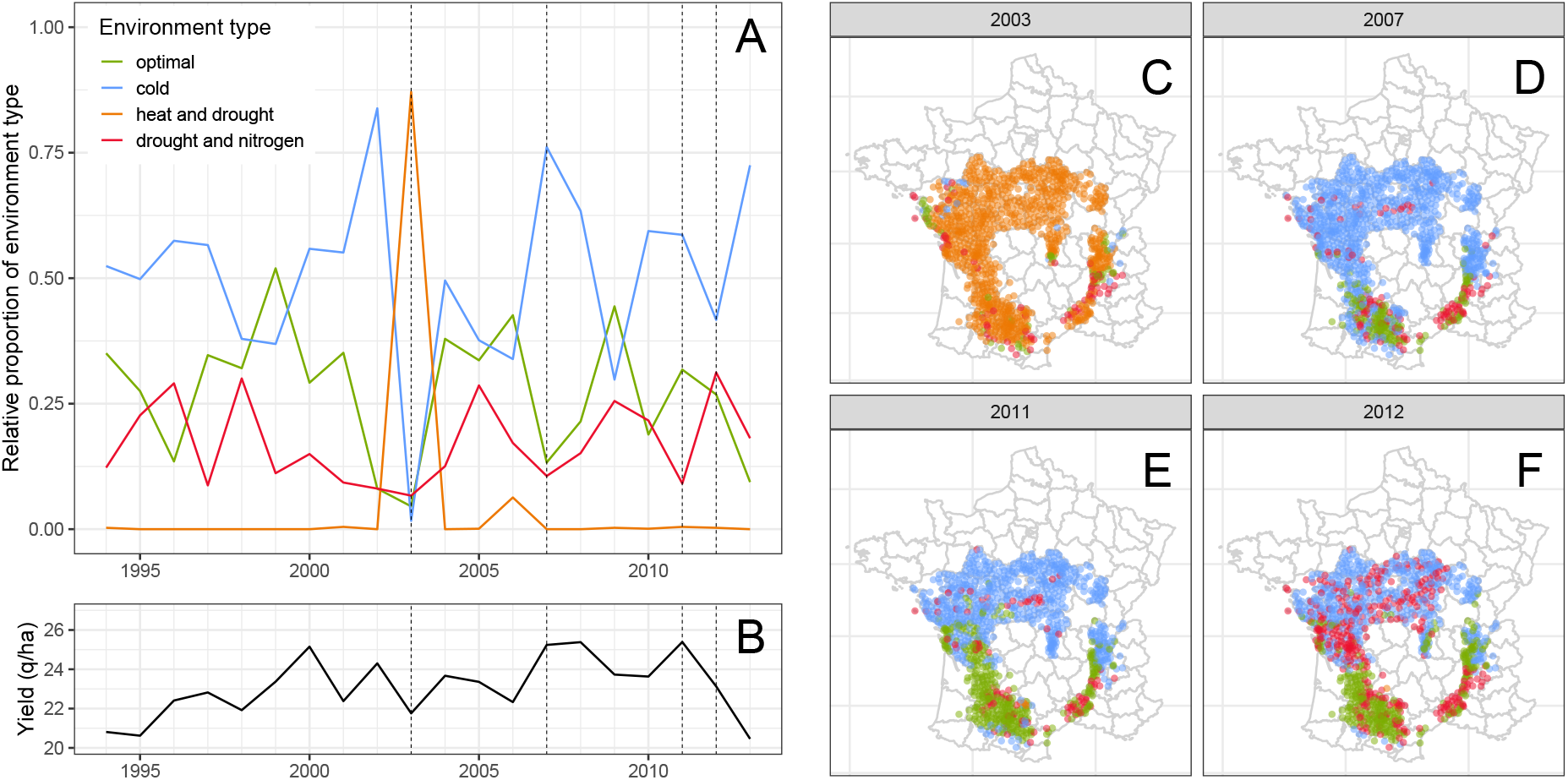
Temporal and spatial distribution of environment types. Panel A displays the evolution of the relative proportion of environment types over 20 years. For reference, panel B is the national sunflower yield. The right panels (C-F) display the spatial distribution of environment types for each individual cropping condition, for a subset of four contrasted years.

The second cluster analysis allowed to group locations with a similar frequency of occurrence of environment types, i.e. locations with predictable abiotic stress patterns. We found that using four groups in the cluster analysis (between:total sum of squares of 86.9 %) yielded contrasted agricultural zones with balanced proportions: from locations with consistently *optimal* environment types (South-West, Figure 6, *Z2*) to consistently *drought and nitrogen* deficit environment types (West, South-East, Figure 6, *Z3*). Still, about 30% of the locations were characterized by an unstable pattern of environment types within the years (labeled with *Z1*, Figure 6).

**Figure 6.**
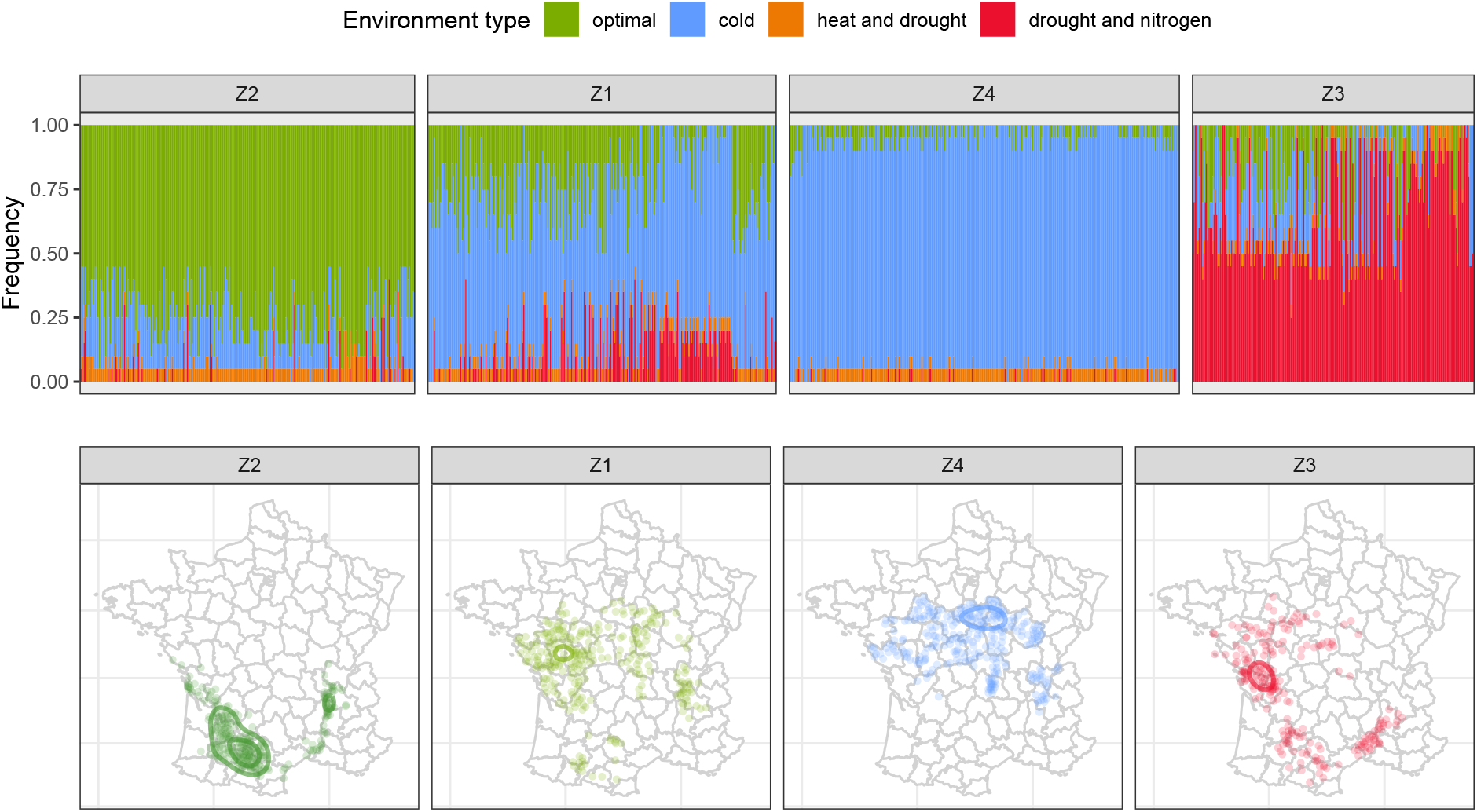
Identification of agricultural zones with a similar frequency of occurrence of environment types. Upper panels represent the frequency of environment types over 20 years (using stacked bars) for each farming location (ordered by increasing longitude). Hierarchical clustering was applied to the frequency of environment types per location, in order to identify groups of locations with a similar frequency of environment types, referred to as agricultural zones (Z1-Z4). In lower panels, the identified agricultural zones are mapped to visualize which geographical positions are sharing common and predictable abiotic stress patterns over the years.

### Optimization of cultivar choice

Using the most frequently sown cultivars in France during the studied period (2009-2013) as a baseline to represent the actual deployment strategy (Figure 7, *reference* strategy), we showed that it was a much better strategy than randomly choosing a cultivar (among the 38 studied and available for sale) for each environment, where yield expectation was lowered by 5.7% (Table 4).

**Figure 7.**
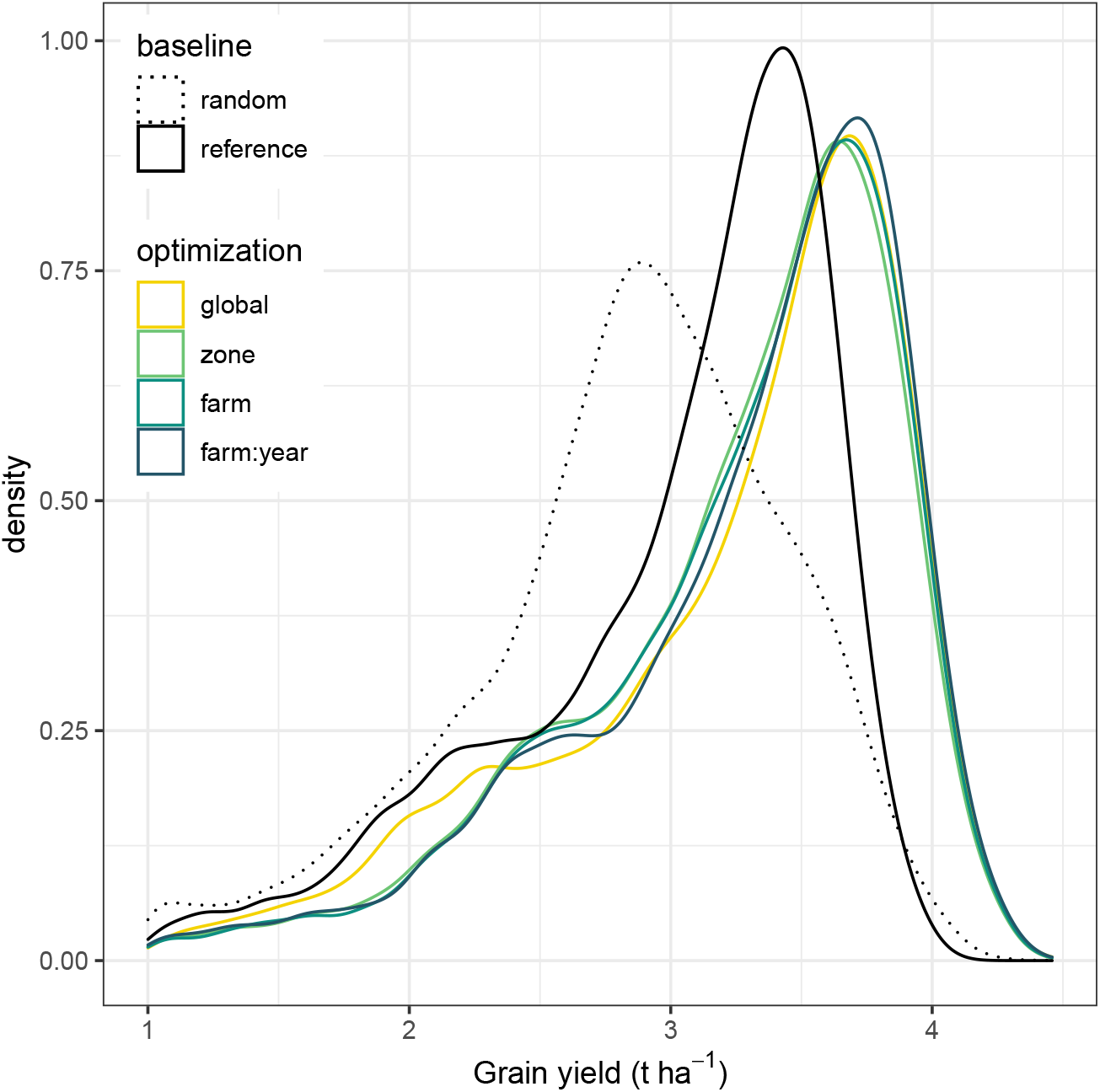
Impact of different cultivar deployment strategies on the distribution of grain yield across the population of cropping conditions. Two baseline deployment strategies are presented: *random*, a random choice of cultivar per farm × year combination (20566 choices), and *reference*, the mean of the most frequently sown cultivars during the studied period. The optimized deployment strategies differed by the number of cultivar choices made across the population of cropping conditions: *global*, one decision for all conditions; *zone*, one decision per agricultural zone (4 choices); *farm*, one decision per farm (1075 choices) and *farm:year*, one decision per farm × year combination (20566 choices).

Figure 7 shows that the grain yield distribution was shifted towards higher yields when changing the deployment strategy from the reference to optimized cultivar choices, whether it was by choosing a single cultivar across all the population of environments or fine-tuning cultivar choice according to yearly cropping conditions on the farm. While the difference among yield distributions was clearly visible between the *reference* and optimized strategies, differences within optimized strategies were more tenuous and occurred mainly for low yield levels.

Yield gain was defined as the relative difference between the considered strategy and reference one. Over the population of environments, the median gain ranged from 7% for the *global* strategy to 8.5% for the *farm:year* strategy (Table 4). The major improvement in overall crop performance was observed between the reference and global strategy, while other strategies brought additional but more substantial gains, with the greatest difference (0.57%) between *year* and *farm:year* strategies. The expected shortfall, a risk measure focusing on the less profitable outcomes, increased more strongly than median gains with more precise deployment strategies. At the farm scale, the yield stability over the 20 sampled years was also increased by 6 - 9.4% when the cultivar choice was optimized.

Finally, we analyzed the yield gain distribution stratified by environment types to further understand how the deployment strategies were differing (Figure 8).

**Figure 8.**
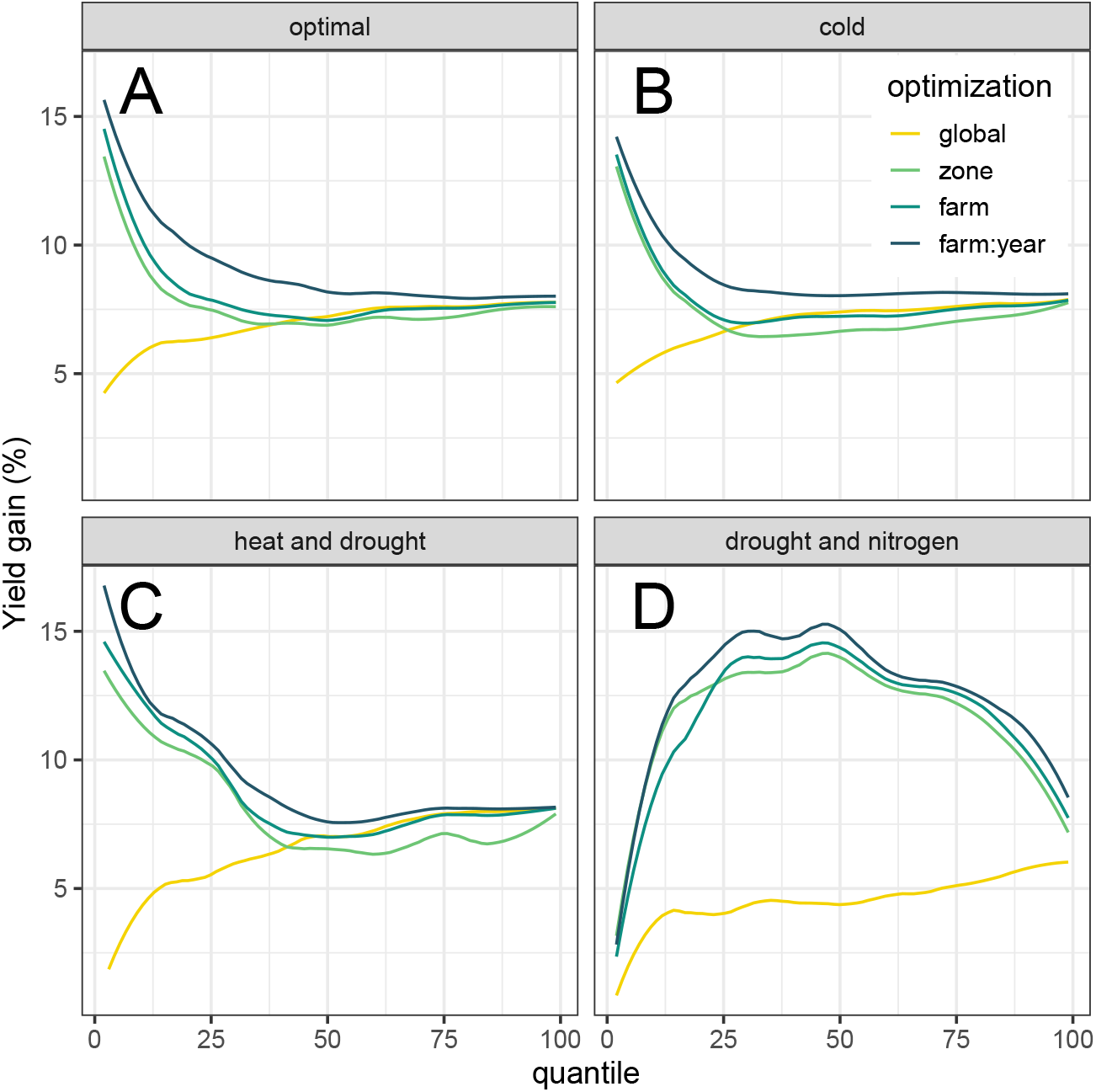
Gain achieved for different cultivar deployment strategies as a function of yield level. For each deployment strategy, the yield gain was computed as the relative difference between a given yield percentile for the tested strategy and the same percentile for the reference strategy. Deployment strategies involving an increased number of cultivar choices are represented with darker shades. The global yield gain distribution was stratified by environment-type and presented in different panels.

We found that moving from a single cultivar choice to specific recommendations had a stronger positive effect when applied to the less productive situations (lowest quantiles). For all environment-types, gains up to 15% can occur. However, with higher yield potential (highest quantiles), the gains allowed by specific recommendations are reduced to match those allowed by the *global* strategy. For the most unfavorable environment-type (*drought and nitrogen*, Figure 8), gains permitted by specific recommendations were higher than with a single cultivar choice, for all the yield quantiles.

## Discussion

We proposed a methodology relying on crop modeling, simulation, and optimization to predict phenotypic plasticity and to decide which cultivar to grow, given environmental conditions and predicted crop performance. This methodology was applied to the sunflower crop at a countrywide scale to assess whether targeted recommendations of cultivars could be considered as a lever to improve crop performance and stability. Working from farmer survey data, we described a large range of climate, soil, and management practices representing various cropping conditions over the country. We then used simulation to extend this initial sampling with a numerical experimental design featuring new cultivars and climates. Computational experiments showed a potential for local recommendations, with gains increasing with the knowledge of pedo-climatic conditions. But the sole pedo-climatic context was not enough to inform the cultivar choice, yield gains were increased when decisions could be made based on environment types, i.e. when diverse geographical locations and years yet experiencing similar abiotic stress patterns were grouped.

The proposed recommendations are dependent not only on the soil and climate data used to build the numerical experiment but also on the methods and model used for the simulation and optimization step. When using two sets of climate data, differing by the downscaling method used in gridded data, we found that the results were robust to these variations. The ranking of deployment strategies was not impacted and the differences between strategies with different climate datasets were 0.45 % maximum. As the crop model directly drives the phenotypic prediction, its ability to capture cultivar differences in response to environment and management drivers is essential. The algorithm of the crop model was evaluated on agricultural extension trials corresponding to a subset of the population of cropping environments simulated in the numerical experiment (Casadebaig et al., 2016). As reviewed by Wang et al. (2019), further progress in the functional modeling of phenotypic plasticity and therefore prediction accuracy is related to the extension of systems biology to plant and crop scale (Hammer et al., 2004; Keurentjes et al., 2011; Poorter et al., 2013). Lastly, the cultivar recommendations based on environment types depend on the methodology used to group similar environments. In this study, we summarized four abiotic drivers over the whole crop cycle, thereby integrating multiple abiotic stresses but missing phenological effects. In Australia, the clustering method used by Chenu et al. (2013) emphasized phenological effects with several sampling points over the crop cycle, by using a moving average on the simulated time series but focused only on water deficit. In France, Beillouin et al. (2018) did not use a simulation model and acted in two steps: (1) fit a regression model (partial least square regression) to select climatic factors that best-explained yield variability and (2) then use the weighted coordinates of the selected model to group environments according to these characteristics. Their approach emphasized the selection of factors that were impacting yield on their dataset. Another option is the clustering method used in Picheny et al. (2017b), where the distance between environments (years) was computed on the complete time series using a non-Euclidian method (dynamic time warping). While the method was applied on raw climatic variables rather than simulated abiotic time series it would take better account of phenology for clustering environments.

### Tactical agronomy

Choosing a cultivar far before sowing is a demanding problem because there is no reliable seasonal weather forecasting for the incoming cropping season on this date. The *global, zone*, and *farm* strategies accounted for this constraint and were based on information available at the sowing date, i.e. the environmental zone or the geographical location. However, the *farm:year* cultivar deployment strategy operates as if a perfect forecast of the incoming climate was available to choose the cultivar. While unrealistic, this strategy allowed to assess the gains expected from a perfect fit of a cultivar and an environment and revealed that this was the only strategy strongly reducing the probability of low yield outcomes at the population level (Figure 8A) and farm level (Table 4).

Another strong assumption of our method is that we do not account for the effect of the distribution system on the actual cultivars made available for each farmer. In France, farmers essentially buy seeds through agricultural cooperative groups, selling different seeds in different regions. In a way, at the farm scale, access to the full genetic diversity is thus reduced, but this distribution system also filters adapted cultivars. On this subject, Sadras and Denison (2016) highlighted how optimization approaches for decision making in crop management are constrained by various trade-offs ultimately impacting the expected gain from these approaches. They suggest that results from optimization methods are particularly useful as null hypothesis, i.e. that the divergence between actual and optimal solutions can help to identify constraints that are relevant to further explore. We could assume that spatial deployment strategies could be nevertheless promoted because national cultivar evaluation is a common feature in Europe (Van Waes, 2009). Official cultivars trials are arranged by dedicated institutes for the examination of Value for Cultivation and Use (VCU). These trials result in the production of *Recommended Variety Lists* and recommendations for cultivation are published either as national summaries or as regional bulletins.

We found that choosing a single cultivar was a good strategy, yielding important gains compared to the reference strategy but that was only marginally lower than more demanding strategies. We explain the relevance of this strategy because the chosen cultivar had the best adaptation to the most frequent environment-type. Favourable environments, encompassing the *optimal* and arguably the *cold* environment types, were very frequent among the population of environments (Figure 6). The cultivar responses to the range of environments were estimated by regressing cultivar yield on site mean (Finlay and Wilkinson, 1963), and the optimal cultivar for the global strategy had the best adaptation to favourable environments (i.e. the highest regression slope, which was > 1) compared to other cultivars.

However, specific types of environments supported more diverse cultivar choices (Table 4), up to 29 distinct cultivars out of 37 candidates for the most precise deployment strategy. But while the diversity of recommended cultivars indeed increased, a list of 6 cultivars was sufficient to cover 99% of cases in the population of cropping environments, indicating that very specific recommendation occurs rarely. We could have expected that more recently released cultivars (after 2010) could have been more frequently recommended because of the genetic gain but the proportion of recent cultivars was 0.63 in the candidate list (n=37), 0.55 in the recommended list (n=29), and 0.5 in the 6 most recommended list. When analyzing the phenotypic features of cultivars, we found that the harvest index value, a strong productivity-related trait (potential seed:biomass ratio), was similar in the recommended and candidate list (p = 0.353). The fact that a recent release date and a raw productivity feature (high harvest index) are not essential for cultivars to be recommended seems to indicate that traits suited for drought-prone environments are available, yet not widespread in current commercial cultivars. But commercial hybrids are probably related because breeders use successful parents for several hybrids, reducing the apparent genetic diversity analyzed in this study. Molecular profiles of sunflower hybrids could be used in a cluster analysis to identify groups, in a similar way that we defined environmental groups.

### Synergy with plant breeding

Genetic progress on sunflower was evaluated at 1.3% per year, in the 1970-2000 period (Vear et al., 2003), with variable gain thorough years. We quantified the median gains provided by educated genotypic choices at 7 - 9% depending on the strategy used but even if this improvement does not scale with time, we could reasonably suppose that it is added to genetic progress. While the exploitation of phenotypic plasticity is often discussed with the perspective of breeding new material (Vega and Chapman, 2006; Nicotra et al., 2010; Messina et al., 2011), model-based cultivar deployment is operational on current genetic material. Moreover, cultivar deployment and breeding should act synergistically as breeding could lead to cultivars with distinctive traits that are more adapted to specific environments. This might be reflected by the steady decrease in acreage share of major cultivars observed in surveys, opening the market to more diverse cultivars.

Regardless of the way the breeding process is organized, it would be always more difficult to assess the specific adaptation of a cultivar to a particular cropping condition than to identify a more widely adapted cultivar (Vincourt and Carolo, 2018). We showed that sowing a widely adapted cultivar in all cropping conditions is an acceptable strategy, however, missing a cultivar match for an unusual cropping condition could be viewed as a wasted opportunity. In this sense, participatory plant breeding might help to produce the observations needed to support a local adaptation strategy (Vincourt and Carolo, 2018).

### Adaptations at the farming system scale

The proposed cultivar deployment strategies considered a single cultivar choice for each farmer per year, because growing sole crops is the mainstream practice according to surveys. The positive effect of species or genetic diversity on ecosystem functioning (diversity–stability hypothesis) was evaluated in different systems, from mixtures of grassland species in the plot scale (e.g. Hector et al., 2010) to mixtures of crops at the farm (Paut et al., 2019) or national scale (Renard and Tilman, 2019). At the farm scale, farmers can opt for a bet-hedging strategy to minimize risk, i.e. growing cultivars having different features in distinct fields, or even by mixing cultivars in a single plot (Barot et al., 2017). In this case, adaptation without the full knowledge of the environmental conditions would allow a better crop performance because of the dampening of the abiotic stress effect at the population level (plants or plots). Among various options to transition to biodiversity-based agriculture (Duru et al., 2015), mixing cultivars probably requires a lesser adaptation for farmers and a lesser challenge for breeders.

## Conclusions

Agricultural surveys were used to describe the population of environments for the sunflower crop in France. Simulation-based optimization revealed a potential for local recommendations, with gains gradually increasing with knowledge of cropping conditions, especially when distinct pedo-climatic conditions were clustered into homogeneous types of environments. At the national scale, tuning the choice of cultivar impacted crop performance the same magnitude as the effect of yearly genetic progress made by breeding. Our results suggest that optimizing the choice of cultivars according to environmental cues deserves consideration as a way to raise and stabilize crop yield at the national and farm scale.

## Acknowledgements

The authors are grateful to the students (Claire Barbet-Massin, Ewen Gery, Bertrand Haquin) and staff from *INRAE* (Céline Colombet, Didier Rafaillac, Colette Quinquiry), *ENSAT* (Michel Labarrère, Pierre Maury) and *Terres Inovia* (Frédéric Bardy, Philippe Christante, André Estragnat, Pascal Fauvin, Céline Motard, Jean-Pierre Palleau, Frédéric Salvi) that helped to constitute the phenotypic database. We also wish to thank the INRA RECORD team (Hélène Raynal, Eric Caselas) for modeling and simulation, and the INRA AgroClim team (Benoît Persyn, Patrick Bertuzzi) that provided gridded climate datasets.

## Funding

Grants were provided by the French Ministry of Agriculture (CASDAR C-2016-03 CARAVAGE) and the French Ministry of Research (ANR SUNRISE ANR-11-BTBR-0005). This work was also supported by the French National Research Agency under the Investments for the Future Program, referred as ANR-16-CONV-0004.

## Competing Interests

The authors have no relevant financial or non-financial interests to disclose. On behalf of all authors, the corresponding author states that there is no conflict of interest.

## Author Contributions

PC, NL, RT, and PD designed and planned the study; PC, AL, EM, and JS provided data; PC, AG, JS, and RT analyzed the data; PC wrote first draft of the manuscript and all authors commented on previous versions of the manuscript. All authors read and approved the final manuscript.

## Data Availability

The datasets generated during and/or analysed during the current study are available from the corresponding author on reasonable request.

## Notes

### Competing Interest Statement

The authors have declared no competing interest.

### Summary of Updates

post-print version

